# An advanced lentil backcross population developed from a cross between *Lens culinaris* × *L. ervoides* for future disease resistance and genomic studies

**DOI:** 10.1101/2021.01.13.426580

**Authors:** Tadesse S Gela, Stanley Adobor, Hamid Khazaei, Albert Vandenberg

## Abstract

Genetically accessible variation to some of the abiotic and biotic stresses are limited in the cultivated lentil (*Lens culinaris* Medik.) germplasm. Introgression of novel alleles from its wild relative species might be required for enhancing the genetic improvement of the crop. *L. ervoides*, one of the wild relatives of lentil, is a proven source of disease resistance for the crop. Here we introduce a lentil advanced backcross population (LABC-01) developed in cultivar CDC Redberry background, based on *L. ervoides* alleles derived from an interspecific recombinant inbred population, LR-59-81. Two-hundred and seventeen individuals of the LABC-01 population at BC_2_F_3:4_ generation were screened for the race 0 of anthracnose (*Colletotrichum lentis*) and stemphylium blight (*Stemphylium botryosum*) under controlled conditions. The population showed significant variations for both diseases and transfer of resistance alleles into the elite cultivar was evident. It also segregated for other traits such as days to flowering, seed coat colour, seed coat pattern and flower colour. Overall, we showed that LABC-01 population can be used in breeding programs worldwide to improve disease resistance and will be available as a valuable genetic resource for future genetic analysis of desired loci introgressed from *L. ervoides*.

## Introduction

Cultivated lentil (*Lens culinaris* Medik.) is an economically important cool-season grain legume with genome size of ∼4.3 Gbp in the haploid complement (Bett *et al*., 2016). The crop is cultivated in more than 70 countries worldwide and the production (2015-2019 average) from Canada (40%), India (19%) and Australia (9%) provides most of the world’s supply. Average global annual production was around 6.3 Mt (FAOSTAT, 2021). A genetic bottleneck limiting genetic variability exists in cultivated lentil germplasm (Erskine *et al*., 1994, 1989; Gupta and Sharma, 2006; Khazaei *et al*., 2016). Broadening the genetic base of breeding programs by introducing new genetic resources is required for development of improved lentil germplasm. Crop wild relatives have been used to improve the resistance and resilience of elite cultivars against various biotic and abiotic stresses in many crops (Hajjar and Hodgkin, 2007) including grain legumes (Pratap *et al*., 2021).

The *Lens* genus has seven closely related taxa; *L. culinaris, L. orientalis* (Boiss.) Hand.-Maz., *L. tomentosus* Ladiz. (primary gene pool); *L. odemensis* Ladiz., *L. lamottei* Czefr. (secondary gene pool); *L. ervoides* (Brign.) Grand. (tertiary gene pool); and *L. nigricans* (M.Bieb.) Grand. (quaternary gene pool, *see* Wong *et al*., 2015). Among them, *L. ervoides* has been identified as a potential source of desirable genes for resistance to major lentil diseases such as anthracnose (*Colletotrichum lentis* Damm) (Tullu *et al*., 2006; Vail *et al*., 2012), ascochyta blight (*Ascochyta lentis* Vassilievsky) (Tullu *et al*., 2010), stemphylium blight (*Stemphylium botryosum* Wallr.) (Podder *et al*., 2013), and fusarium wilt (*Fusarium oxysporum* f. sp. *lentis*) (Singh *et al*., 2017) along with yield and its components (Gupta and Sharma, 2006; Tullu *et al*., 2011; Tullu *et al*., 2013; Chen, 2018), and abiotic stresses (Gorim and Vandenberg, 2017; Yuan *et al*., 2017). Introgression of the desirable alleles from *L. ervoides* to *L. culinaris* elite germplasm were facilitated by embryo/ovule rescue techniques that overcome the interspecific reproductive barriers (Fiala *et al*., 2009; Tullu *et al*., 2013). However, the introgressed gene from a distant wild relative into elite cultivars may result in disruption of the long-accumulated agronomic and quality traits due to linkage drag and/or epistatic interactions of deleterious genes of undesired wild traits (Tanksley *et al*., 1989; Tanksley and Nelson, 1996). In many cases, these undesired traits are dominant and polygenic, making it difficult to select against and impeding the interspecific hybrid progeny from direct use in the breeding programs. In lentil, Tullu *et al*. (2013) and Chen (2018) reported the presence of undesired traits such as seed dormancy, poor emergence, extremely small seed size, and pod dehiscence in *L. ervoides* interspecific lines.

The advanced backcross (AB) populations are developed through multiple backcrossing (BC_2_ or BC_3_) followed by multiple rounds of selfing, and they may contain single or multiple, fixed, or non-fixed segments of the introgressed genome of the wild species (Fulton *et al*., 1997). The AB lines are useful genetic materials for the development of introgression lines, which consist of fixed lines that are carrying a single or a few genomic segments associated with desired traits (Frischa *et al*., 1999, Prohens *et al*., 2017, Dempewolf *et al*., 2017). To make the introgression process more efficient and applicable, Tanksley and Nelson (1996) proposed an advanced backcross-quantitative trait loci (AB-QTL) mapping approach as a tool to minimize the undesirable segments of the wild genome through repeated backcrossing to the elite cultivar and simultaneous mapping of QTL underlying the trait of interest. The AB-QTL strategy has been used in many crop species for identifying introgression QTL for many traits of interest (*reviewed by* Bhanu *et al*., 2017) including disease resistance (Yun *et al*., 2006; Schmalenbach *et al*., 2008; Taguchi-Shiobara *et al*., 2013).

Here we introduce a lentil advanced backcross population (LABC-01) derived from a cross between the CDC Redberry and an interspecific line, LR-59-81 developed from *L. ervoides*. This population offers an opportunity to utilize beneficial traits introgressed from lentil wild relatives. As a showcase, the responses of the LABC-01 population at BC_2_F_3:4_ generation to anthracnose race 0 and stemphylium blight under climate-controlled conditions are presented.

## Experimental details

### Parental lines selection

A lentil advanced backcross population (LABC-01) was developed from two founder lines, CDC Redberry and LR-59-81 (Figure 1A). The recurrent parent, CDC Redberry, was a red lentil cultivar released by the Crop Development Center (CDC), University of Saskatchewan (USask) for its high yield and partial resistance to anthracnose race 1 and ascochyta blight (Vandenberg *et al*., 2006). CDC Redberry was used to develop a lentil reference genome (Bett *et al*., 2016). The line LR-59-81 was selected from the LR-59 interspecific recombinant inbred line (RIL) population (Fiala *et al*., 2009), which was developed from a cross between *L. culinaris* cv. Eston × *L. ervoides* accession L-01-827A (Figure 1B). Embryo rescue techniques were used to obtain the F_1_ seeds (Fiala *et al*., 2009). Line LR-59-81 was evaluated for resistance to anthracnose (race 0 and 1), ascochyta blight and stemphylium blight (Table 1) and has been commonly used as a source of resistant for both races of anthracnose (Banniza *et al*., 2018; Gela *et al*., 2020).

**Table 1.** Disease response and morphological characteristics of parental lines of the LABC-01 population.

| Characteristics | CDC Redberry | LR-59-81/L-01-827A | Reference |
| --- | --- | --- | --- |
| Disease resistance |  |  |  |
| Anthracnose race 0 | Susceptible | Resistant | Vail *et al.* (2012), Gela *et al.* (2020) |
| Anthracnose race 1 | Partially Resistant | Resistant | Banniza *et al.* (2018) |
| Ascochyta blight | Resistant | Resistant | Tullu *et al.* (2010) |
| Stemphylium blight | Susceptible | Resistant | Podder *et al.* (2013) |
| Days to flower | Late | Early | This study |
| Days to maturity | Late | Early | Chen (2018) |
| Seed cotyledon colour | Red | Red | This study |
| Seed coat color/pattern | Gray/unpatterned | Brown/black marble | This study |
| Seed weight | High | Low | Chen (2018) |
| Flower colour | White with light blue veins | Purple | This study |
| Plant height | Tall | Short/weak stem | Chen (2018) |

**Figure 1.**
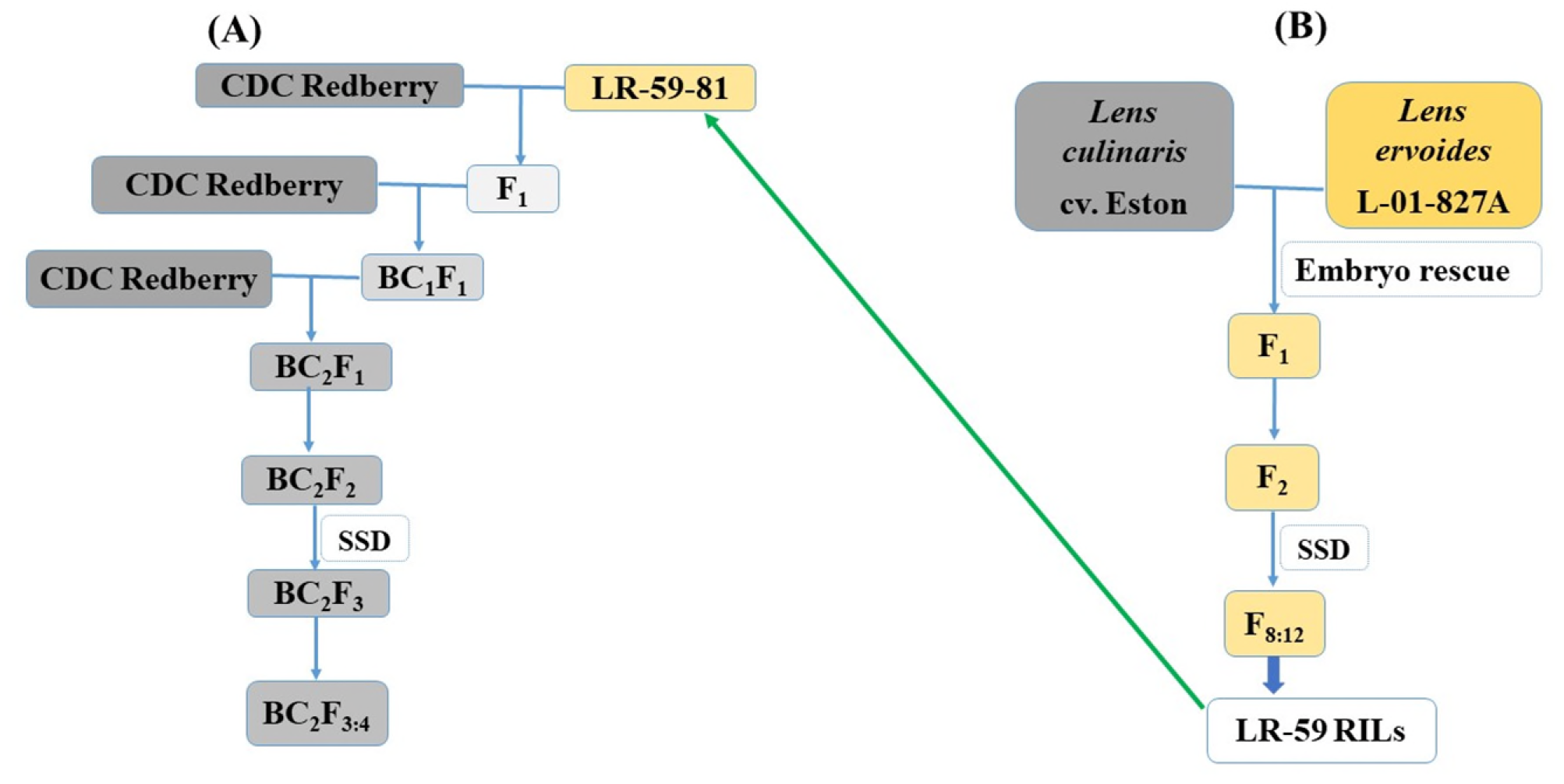
Schematic diagram of lentil advanced backcross mapping population (LABC-01) development (**A**) and the pedigree of the LR-59-81 (**B**). SSD, single seed descent.

### LABC-01 population development

The LABC-01 population was developed by crossing CDC Redberry × LR-59-81 to obtain the F_1_ generation (Figure 1A). All F_1_ seeds were fertile and hybridity of F_1_ plants was confirmed by flower colour as a morphological marker. CDC Redberry had typical *L. culinaris* flower-type, white background with light blue veins and LR-59-81 had purple flowers (typical *L. ervoides*). All F_1_ plants had purple flowers. Purple flower colour was dominant over white with blue veins and it is known to be inherited as a simple Mendelian fashion (Singh *et al*., 2014). Two F_1_ plants were backcrossed to CDC Redberry to obtain BC_1_F_1_ seeds. To avoid genetic drift, all efforts were made to achieve the maximum number of cross combinations. A total of 111 and 73 BC_1_F_1_ seeds were harvested from two F_1_ plants, respectively. The segregation of the flower colours was checked for the BC_1_F_1_ population and fit a 1:1 ratio (88 white: 93 purple, chi-square (χ^2^) _(1:1)_ = 0.138, *P* = 0.710), indicating unbiased segregation of the BC_1_F_1_. The second backcross was made independently with all 184 BC_1_F_1_ plants to generate the BC_2_ population, and one to two BC_2_F_1_ seeds were advanced to BC_2_F_2_ for each successful BC_1_F_1_ backcross. A total of 217 BC_2_F_2_ individuals was generated and one seed of each individual was arbitrarily selected and selfed to generate the BC_2_F_3_ generation and onward using a single seed descent approach.

### Growth conditions

In all experiments, the growth chamber conditions were adjusted to 18 h light and 6 h dark, with the temperatures maintained at 21 °C (day) and 18 °C (night) and the photosynthesis photon flux density was set to 300 μmol m^-2^ s^-1^ during the light period at the crop canopy level. All experiments were carried out at the controlled-climate growth chambers at USask College of Agriculture and Bioresources phytotron facility, Saskatoon, Canada.

### Disease phenotyping

#### Phenotyping for anthracnose resistance

A total of 217 BC_2_F_3:4_ individuals of the LABC-01 population and parental lines were evaluated for anthracnose race 0 under growth chamber conditions. Fungal inoculum production, inoculation and plant growth conditions were performed as described by Gela *et al*. (2020). Briefly, two plants of each line were grown in a set of 38-cell cone trays (26.8 × 53.5 cm) per replication filled with SUNSHINE Mix #4 plant growth medium (Sun Gro Horticulture, Seba Beach, AB, Canada) and perlite (Specialty Vermiculite Canada, Winnipeg, MB) in a 3:1 ratio (v/v). The susceptible control cv. Eston (Slinkard, 1981) and the parental lines were included in each tray. The experiment design was a randomized complete block (RCBD) with seven replicates. Replicates were inoculated over time. Four-week-old seedlings were inoculated with a spore suspension (5 × 10^4^ spores mL^-1^) of *C. lentis* race 0 isolate CT-30 (Banniza *et al*., 2018) at 3 mL per plant using an airbrush. Individual plants were scored for anthracnose severity at 8-10 days post inoculation (dpi) using a 0 to 10 rating scale with 10% increments. The final disease score from each plant was the mean score per replicate and the data were converted to percent disease severity using the class midpoints for statistical analysis.

#### Phenotyping for stemphylium blight resistance

Six seeds of each individual line of the LABC-01 population were sowed in a 10-cm plastic pots filled with SUNSHINE Mix #4 plant growth medium and arranged in RCBD with three replicates. Two weeks post emergence, plants were thinned to four plants per replicate and fertilized once every week using 3 gL^-1^ of soluble N:P:K (20:20:20) PlantProd^®^ fertilizer (Nu-Gro Inc., Brantford, ON, Canada). Cultivars Eston (Slinkard, 1981) and CDC Glamis (Vandenberg *et al*., 2002) were used as resistant and susceptible checks, respectively.

A culture stock of the aggressive *S. botryosum* SB19 isolate collected from Southeast Saskatchewan was obtained from the Plant Pathology Laboratory, USask for mass spore production following a procedure described by Caudillo-Ruiz (2016). Plants were spray-inoculated at the pre-flowering stage with approximately 3 mL of conidial suspension per plant at a concentration of 1 × 10^5^ conidia mL^-1^ using an airbrush (Badger Airbrush, model TC 20, USA). Two droplets of Tween^®^ 20 (Sigma, Saint Louis, MO, USA) were added to every 1000 mL of suspension before inoculation to help reduce the surface tension of water and promote plant tissue contact. Plants were placed in an incubation chamber for seven days. Two humidifiers (Vicks Fabrique Paz Canada, Inc., Milton, ON, Canada) were placed in the incubation chamber to ensure 90-100% relative humidity for infection and disease development. Blocks were inoculated over time.

Disease severity was assessed visually at 7 dpi using a semi-quantitative rating scale (0-10) where 0, healthy plants; 1, few tiny lesions; 2, a few chlorotic lesions; 3, expanding lesions on leaves, onset of leaf drop; 4, 1/5th of nodes affected by lesions and leaf drop; 5, 2/5th of nodes affected; 6, 3/5th of nodes affected; 7, 4/5th of nodes affected; 8, all leaves dried up; 9, all leaves dried up but stem green; and 10, plant completely dead. Disease severity was assessed on single plants within the experimental unit (pot). For each genotype, four single plants per replicate pot were assessed. Disease severity data was analyzed using the median disease severity score for each genotype.

### Data analyses

Statistical analyses were conducted for both anthracnose and stemphylium blight severity using SAS software (SAS 9.4, SAS Institute, Cary, North Carolina, 2011). Normality and variance homogeneity of the residuals were tested using the Shapiro-Wilk normality test and Levene’s test for homogeneity, respectively. The data did not fit the assumptions of a Gaussian distribution and were normalised using a lognormal distribution in the GLIMMIX procedure. Genotypes were treated as fixed effect and blocks as random effect and significance of variances were declared at 5% significance level. Least square means were estimated for genotype using LSMEANS statements.

## Results and discussion

In this study we developed a lentil advanced backcross population to explore the valuable genetic variation introgressed from lentil wild relative *L. ervoides* into adapted cultivar CDC Redberry. *L. ervoides* accession L-01-827A, the parent to the interspecific LR-59-81, has previously shown adaptation to drought (Gorim and Vandenberg, 2017) and resistance to diseases such as ascochyta blight (Tullu *et al*., 2010), stemphylium blight (Podder *et al*., 2013), anthracnose (Vail *et al*., 2012; Gela *et al*., 2020), and the parasitic weed broomrape (*Orobanche crenata* Forsk.) (Bucak *et al*., 2014). Our results revealed variation for desirable traits in the LABC-01 population that inherited from L-01-827A in the cultivated lentil background including disease resistance and phenological traits.

The LABC-01 population could possibly combine important key traits from *L. ervoides* for lentil genetic improvement as a pre-breeding genetic source and as a valuable resource on which to conduct further genetic studies. Our data showed that BC_2_F_3:4_ generation had a continuous distribution for days to flowering (Supplemental Figure S1) and also segregated for morphological traits such as flower colour (191 white: 26 purple), seed coat colour (190 gray: 27 tan), and seed coat pattern (185 absent: 32 marbled). Vail (2010) and Chen (2018) have reported segregation of several agronomic and phenotypic characteristics including plant vigour, yield and its components in the genetic populations derived from LR-59-81 or L-01-827A. A similar trend was also reported for seed iron concentration (Podder, 2018). Multi-location evaluation of LABC-01 lines will be considered necessary for genetic analysis of these traits and selection of advanced lines for the lentil breeding programs.

Significant variation for anthracnose race 0 resistance was observed among the 217 LABC-01 individuals (F-value= 3.98, *P*<0.0001). The LR-59-81 had a resistant reaction with a mean disease severity of 36%, whereas the recurrent parent CDC Redberry showed susceptible reactions with a mean of 85%. A large number of the LABC-01 individuals showed to be susceptible to race 0. The disease severity ranged from 17-95% with a mean of 70.2%. Transgressive variation for race 0 resistance relative to that of the resistant LR-59-81 was observed (Figure 2A) which is consistent with the findings (Fiala *et al*., 2009; Tullu *et al*., 2013), who reported the transgressive segregation and skewed distributions of lines toward the higher level of disease severity. Two pathogenic races of anthracnose (races 0 and 1) have been described (Buchwaldt *et al*., 2004). Resistance to race 1 is abundant in cultivated lentil germplasm, however, resistance to the more virulent race 0 is mainly limited to *L. ervoides* (Tullu *et al*., 2006; Barilli *et al*., 2020; Gela *et al*., 2020). Transfer of race 0 resistance alleles into the cultivated background could widen the lentil breeding genepool and provide benefits for cultivar development.

**Figure 2.**
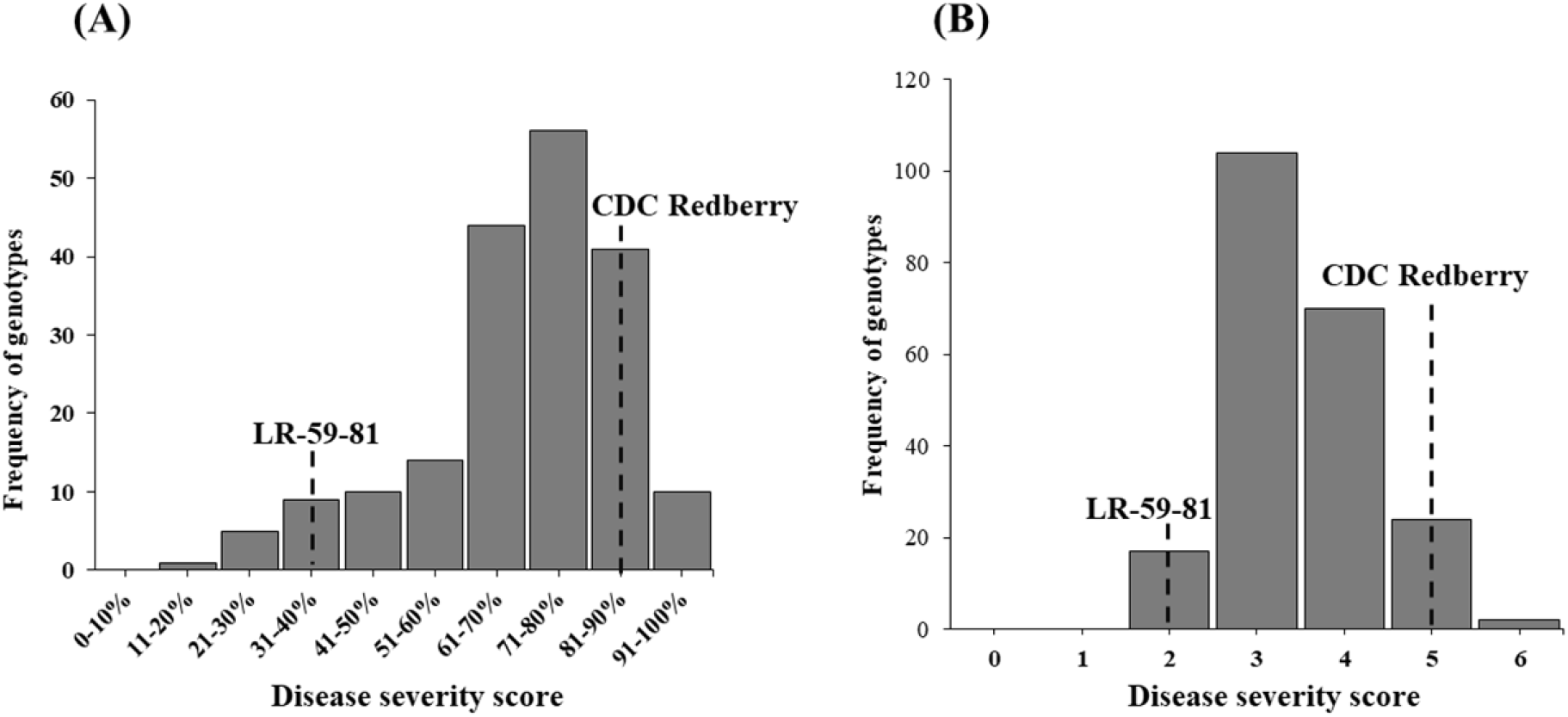
Frequency distribution of mean anthracnose race 0 (n = 217, *P(W)* = 0.001) (**A**) and stemphylium blight severity (n = 217, *P(W)* = 0.018) (**B**) for LABC-01 population at BC_2_F_3:4_ generation under growth chamber condition. The vertical lines indicate the average disease severity of the parents. *P (W)*, P value of the Shapiro–Wilk test for normality.

Significant variation in stemphylium blight severity was observed among the 217 LABC-01 individuals (F-value= 1.81, *P*<0.0001). The resistant line, LR-59-81, had significantly less disease severity (2.32) compared to the resistant check cv. Eston (4.16) and recipient parent CDC Redberry (5.13). The distribution of disease severity as a measure of stemphylium blight response for LABC-01 lines showed continuous variation ranging from 1.59 to 5.80 (Figure 2B), suggesting polygenic regulation of stemphylium blight severity. None of the LABC-01 individuals had significantly less disease than the resistant parent LR-59-81 (Supplementary Table S1). This observation is consistent with the findings of Adobor *et al*. (2020) who also reported the absence of resistant transgressive segregants in *L. ervoides* interspecific population screened for stemphylium blight resistance in the greenhouse, growth chamber and the field conditions. A high proportion of the LABC-01 individuals (144) had similar disease severity when compared to the resistant parental line LR-59-81, indicating that resistance genes were transferred from the resistant parent to LABC-01 individual lines (Supplementary Table S1).

Understanding the genetic architecture of the favourable traits from the wild germplasm provides breeders information that can aid in the introgression of the traits while avoiding linkage drag of deleterious characteristics of the wild species (Tanksley and Nelson, 1996). The LABC-01 population can be used for preliminary QTL mapping and genetic characterization of agronomic traits and disease resistance that have been introgressed into CDC Redberry. Since no selection was carried out during population creation, some of the LABC-01 lines may be of interest as starting materials for the development of fixed introgression populations for specific traits of interest (Prohens *et al*., 2017). For instance, QTL analysis can be conducted with the anthracnose race 0 and stemphylium blight data and then the identified markers can be used to facilitate the development of introgression lines (ILs) such chromosome segment substitution lines (CSSLs) and/or near isogenic lines (NILs) by means of marker-assisted selection. The ILs are important for fine QTL mapping studies and developing genetically characterized elite materials that can be directly incorporated into breeding programs (Zamir, 2001; Eduardo *et al*., 2005, Tian *et al*., 2006). The present population is being genotyped using an exome capture array described by Ogutcen *et al*. (2018) which will provide a valuable genetic tool for collaborative lentil research. The plant materials are currently managed and stored at the Crop Development Centre, University of Saskatchewan, Saskatoon, Canada.

## Acknowledgements

The authors gratefully acknowledge funding from the Natural Sciences and Engineering Research Council of Canada (NSERC) Industrial Research Chair Program, the Saskatchewan Pulse Growers, and the University of Saskatchewan. We are also thankful for the technical assistance of the Pulse Pathology lab staff and Pulse Crop Breeding and Genetics group at the University of Saskatchewan.

