## Supplementary Table S1 for "An advanced lentil backcross population developed from a cross between *Lens culinaris* × *L. ervoides* for future disease resistance and genomic studies"

**Supplementary Table S1.** Mean anthracnose race 0 (A0) and stemphylium blight (SB) severity for 217 LABC-01 population and comparison with the resistance of LR-59-81 donor parental line screened under growth chamber conditions. The genotypes marked with bold font are the top resistant lines to anthracnose race 0 that also have resistance to stemphylium blight comparable to LR-59-81.

| Entry | Genotype | A0 | SEM | Sig. | SB | SEM | Sig. |
|---|---|---|---|---|---|---|---|
| 1 | LABC-01-1 | 83.2 | 0.13 | ** | 2.71 | 0.54 | ns |
| 2 | LABC-01-2 | 70.22 | 0.11 | ** | 2.41 | 0.48 | ns |
| 3 | LABC-01-3 | 82.34 | 0.13 | ** | 3.74 | 0.75 | ns |
| 4 | LABC-01-4 | 76.83 | 0.12 | ** | 2.62 | 0.52 | ns |
| 5 | LABC-01-5 | 61.16 | 0.1 | * | 3.56 | 0.71 | ns |
| 6 | LABC-01-6 | 76.83 | 0.12 | ** | 3.11 | 0.62 | ns |
| 7 | LABC-01-7 | 36.53 | 0.06 | ns | 4.16 | 0.83 | * |
| 8 | LABC-01-8 | 69.06 | 0.11 | ** | 3.98 | 0.8 | * |
| 9 | LABC-01-9 | 67.3 | 0.11 | ** | 3.04 | 0.61 | ns |
| 10 | LABC-01-10 | 62.27 | 0.1 | * | 2.88 | 0.58 | ns |
| 11 | LABC-01-11 | 58.58 | 0.09 | * | 3.02 | 0.6 | ns |
| 12 | LABC-01-12 | 57.3 | 0.09 | * | 5.16 | 1.03 | ** |
| 13 | LABC-01-13 | 77.14 | 0.12 | ** | 1.59 | 0.32 | ns |
| 14 | LABC-01-14 | 80.16 | 0.13 | ** | 3.17 | 0.64 | ns |
| 15 | LABC-01-15 | 54.88 | 0.09 | ns | 3.48 | 0.7 | ns |
| 16 | LABC-01-16 | 55.91 | 0.09 | * | 2.29 | 0.46 | ns |
| 17 | LABC-01-17 | 66.6 | 0.11 | ** | 3.11 | 0.62 | ns |
| 18 | LABC-01-18 | 88.38 | 0.14 | *** | 4.04 | 0.81 | * |
| 19 | LABC-01-19 | 59.44 | 0.1 | * | 4.22 | 0.84 | * |
| 20 | LABC-01-20 | 57.19 | 0.09 | * | 3.23 | 0.65 | ns |
| 21 | LABC-01-21 | 72.96 | 0.12 | ** | 3.78 | 0.76 | * |
| 22 | LABC-01-22 | 68.35 | 0.11 | ** | 3.48 | 0.7 | ns |
| 23 | LABC-01-23 | 82.95 | 0.13 | ** | 3.46 | 0.69 | ns |
| 24 | LABC-01-24 | 42.6 | 0.07 | ns | 2.52 | 0.5 | ns |
| 25 | LABC-01-25 | 71.79 | 0.12 | ** | 3 | 0.6 | ns |
| 26 | LABC-01-26 | 61.45 | 0.1 | * | 2.62 | 0.52 | ns |
| 27 | LABC-01-27 | 65.63 | 0.11 | ** | 2.74 | 0.55 | ns |
| 28 | LABC-01-28 | 63.24 | 0.1 | ** | 3.3 | 0.66 | ns |
| 29 | LABC-01-29 | 68.94 | 0.11 | ** | 2.92 | 0.59 | ns |
| 30 | LABC-01-30 | 59.08 | 0.1 | * | 5.31 | 1.06 | ** |
| 31 | LABC-01-31 | 69.51 | 0.11 | ** | 4.83 | 0.97 | ** |

| Entry | Genotype | A0 | SEM | Sig. | SB | SEM | Sig. |
|---|---|---|---|---|---|---|---|
| 32 | LABC-01-32 | 53.96 | 0.09 | ns | 3.62 | 0.72 | ns |
| 33 | LABC-01-33 | 55.2 | 0.09 | * | 4.48 | 0.9 | ** |
| 34 | LABC-01-34 | 61.42 | 0.1 | * | 4.29 | 0.86 | * |
| 35 | LABC-01-35 | 85.93 | 0.14 | *** | 3.56 | 0.71 | ns |
| 36 | LABC-01-36 | 72.9 | 0.12 | ** | 5.09 | 1.02 | ** |
| 37 | LABC-01-37 | 74.55 | 0.12 | ** | 3.39 | 0.68 | ns |
| 38 | LABC-01-38 | 48.74 | 0.08 | ns | 2.73 | 0.55 | ns |
| **39** | **LABC-01-39** | **32.99** | **0.05** | **ns** | **3.11** | **0.62** | **ns** |
| 40 | LABC-01-40 | 57.68 | 0.09 | * | 2.29 | 0.46 | ns |
| 41 | LABC-01-41 | 59.62 | 0.1 | * | 5.8 | 1.16 | ** |
| 42 | LABC-01-42 | 80.59 | 0.13 | ** | 3.3 | 0.66 | ns |
| 43 | LABC-01-43 | 76.96 | 0.12 | ** | 3.11 | 0.62 | ns |
| 44 | LABC-01-44 | 81.52 | 0.13 | ** | 4.82 | 0.96 | ** |
| 45 | LABC-01-45 | 84.16 | 0.14 | *** | 5.13 | 1.03 | ** |
| 46 | LABC-01-46 | 68.13 | 0.11 | ** | 3.74 | 0.75 | ns |
| **47** | **LABC-01-47** | **23.92** | **0.04** | **ns** | **3.42** | **0.68** | **ns** |
| 48 | LABC-01-48 | 71.97 | 0.12 | ** | 3.16 | 0.63 | ns |
| **49** | **LABC-01-49** | **33.47** | **0.05** | **ns** | **2.88** | **0.58** | **ns** |
| 50 | LABC-01-50 | 54.31 | 0.09 | ns | 3.3 | 0.66 | ns |
| 51 | LABC-01-51 | 83.31 | 0.13 | ** | 4.12 | 0.82 | * |
| 52 | LABC-01-52 | 50.27 | 0.08 | ns | 3.56 | 0.71 | ns |
| 53 | LABC-01-53 | 73.7 | 0.12 | ** | 3.3 | 0.66 | ns |
| 54 | LABC-01-54 | 63.54 | 0.1 | ** | 5.3 | 1.06 | ** |
| 55 | LABC-01-55 | 74.83 | 0.12 | ** | 4.12 | 0.82 | * |
| 56 | **LABC-01-56** | **39.81** | **0.06** | **ns** | **3.11** | **0.62** | **ns** |
| 57 | LABC-01-57 | 22.99 | 0.04 | * | 5.3 | 1.06 | ** |
| 58 | LABC-01-58 | 74.47 | 0.12 | ** | 2.62 | 0.52 | ns |
| 59 | LABC-01-59 | 55.45 | 0.09 | * | 3.87 | 0.77 | * |
| 60 | LABC-01-60 | 69.94 | 0.11 | ** | 2.97 | 0.59 | ns |
| 61 | LABC-01-61 | 64.45 | 0.1 | ** | 2.38 | 0.48 | ns |
| 62 | LABC-01-62 | 87.83 | 0.14 | *** | 5.57 | 1.12 | ** |
| 63 | LABC-01-63 | 76.04 | 0.12 | ** | 3.02 | 0.6 | ns |
| 64 | LABC-01-64 | 32.67 | 0.05 | ns | 4.12 | 0.82 | * |
| 65 | LABC-01-65 | 69.68 | 0.11 | ** | 4.81 | 0.96 | ** |
| 66 | LABC-01-66 | 18.86 | 0.03 | ** | 5.38 | 1.08 | ** |
| 67 | LABC-01-67 | 15.54 | 0.02 | ** | 4.33 | 0.87 | * |
| 68 | LABC-01-68 | 61.86 | 0.1 | * | 3.3 | 0.66 | ns |
| 69 | LABC-01-69 | 84.68 | 0.14 | *** | 3.3 | 0.66 | ns |

| Entry | Genotype | A0 | SEM | Sig. | SB | SEM | Sig. |
|---|---|---|---|---|---|---|---|
| 70 | LABC-01-70 | 65.36 | 0.11 | ** | 3.13 | 0.63 | ns |
| 71 | LABC-01-71 | 51.44 | 0.08 | ns | 3.43 | 0.69 | ns |
| 72 | LABC-01-72 | 55.62 | 0.09 | * | 4.76 | 0.95 | ** |
| 73 | LABC-01-73 | 74.19 | 0.12 | ** | 4.44 | 0.89 | ** |
| 74 | LABC-01-74 | 75.4 | 0.12 | ** | 3.7 | 0.74 | ns |
| 75 | LABC-01-75 | 60.33 | 0.1 | * | 4.45 | 0.89 | ** |
| 76 | LABC-01-76 | 59.83 | 0.1 | * | 3.27 | 0.65 | ns |
| 77 | LABC-01-77 | 84.56 | 0.14 | *** | 3.4 | 0.68 | ns |
| 78 | LABC-01-78 | 76.06 | 0.12 | ** | 1.96 | 0.39 | ns |
| 79 | LABC-01-79 | 64.64 | 0.1 | ** | 2.97 | 0.59 | ns |
| 80 | LABC-01-80 | 87 | 0.14 | *** | 4.79 | 0.96 | ** |
| 81 | **LABC-01-81** | **28.61** | **0.05** | **ns** | **3.11** | **0.62** | **ns** |
| 82 | LABC-01-82 | 63.44 | 0.1 | ** | 3.48 | 0.7 | ns |
| 83 | LABC-01-83 | 69.34 | 0.11 | ** | 2.88 | 0.58 | ns |
| 84 | LABC-01-84 | 59.36 | 0.1 | * | 3.68 | 0.74 | ns |
| 85 | LABC-01-85 | 82.38 | 0.13 | ** | 3.3 | 0.66 | ns |
| 86 | LABC-01-86 | 47.26 | 0.08 | ns | 2.82 | 0.56 | ns |
| 87 | LABC-01-87 | 61.61 | 0.1 | * | 3.91 | 0.78 | * |
| 88 | LABC-01-88 | 44.7 | 0.07 | ns | 3.62 | 0.72 | ns |
| 89 | **LABC-01-89** | **40.84** | **0.07** | **ns** | **2.71** | **0.54** | **ns** |
| 90 | LABC-01-90 | 41.87 | 0.07 | ns | 2.82 | 0.56 | ns |
| 91 | LABC-01-91 | 81.95 | 0.13 | ** | 2.29 | 0.46 | ns |
| 92 | LABC-01-92 | 58.43 | 0.09 | * | 4.5 | 0.9 | ** |
| 93 | LABC-01-93 | 47.09 | 0.08 | ns | 1.71 | 0.34 | ns |
| 94 | LABC-01-94 | 46.84 | 0.08 | ns | 3.21 | 0.64 | ns |
| 95 | LABC-01-95 | 63.37 | 0.1 | ** | 3.46 | 0.69 | ns |
| 96 | LABC-01-96 | 58.63 | 0.09 | * | 3.66 | 0.73 | ns |
| 97 | LABC-01-97 | 41.78 | 0.07 | ns | 4.07 | 0.81 | * |
| 98 | LABC-01-98 | 75.89 | 0.12 | ** | 3 | 0.6 | ns |
| 99 | LABC-01-99 | 51.02 | 0.08 | ns | 4.22 | 0.84 | * |
| 100 | LABC-01-100 | 88.85 | 0.14 | *** | 3.62 | 0.72 | ns |
| 101 | LABC-01-101 | 74.68 | 0.12 | ** | 2.19 | 0.44 | ns |
| 102 | LABC-01-102 | 91.54 | 0.15 | *** | 3.11 | 0.62 | ns |
| 103 | **LABC-01-103** | **33.97** | **0.05** | **ns** | **3.23** | **0.65** | **ns** |
| 104 | **LABC-01-104** | **34.88** | **0.06** | **ns** | **2.82** | **0.56** | **ns** |
| 105 | LABC-01-105 | 59.16 | 0.1 | * | 4.31 | 0.86 | * |
| 106 | LABC-01-106 | 54.72 | 0.09 | ns | 4.38 | 0.88 | * |
| 107 | LABC-01-107 | 86.13 | 0.14 | *** | 4.42 | 0.89 | ** |

| Entry | Genotype | A0 | SEM | Sig. | SB | SEM | Sig. |
|---|---|---|---|---|---|---|---|
| 108 | LABC-01-108 | 72.19 | 0.12 | ** | 3.11 | 0.62 | ns |
| 109 | LABC-01-109 | 80.36 | 0.13 | ** | 3.43 | 0.69 | ns |
| 110 | LABC-01-110 | 88.63 | 0.14 | *** | 3.85 | 0.77 | * |
| 111 | LABC-01-111 | 79 | 0.13 | ** | 4.5 | 0.9 | ** |
| 112 | LABC-01-112 | 58.21 | 0.09 | * | 2.62 | 0.52 | ns |
| 113 | LABC-01-113 | 69.01 | 0.11 | ** | 3.11 | 0.62 | ns |
| 114 | LABC-01-114 | 75.42 | 0.12 | ** | 3.3 | 0.66 | ns |
| 115 | LABC-01-115 | 69.05 | 0.11 | ** | 3.43 | 0.69 | ns |
| 116 | LABC-01-116 | 94.15 | 0.15 | *** | 3.56 | 0.71 | ns |
| 117 | LABC-01-117 | 78.82 | 0.13 | ** | 3.23 | 0.65 | ns |
| 118 | LABC-01-118 | 85.73 | 0.14 | *** | 4.48 | 0.9 | ** |
| 119 | LABC-01-119 | 60.65 | 0.1 | * | 2.47 | 0.49 | ns |
| 120 | LABC-01-120 | 44.12 | 0.07 | ns | 3.16 | 0.63 | ns |
| 121 | LABC-01-121 | 72.92 | 0.12 | ** | 2.97 | 0.59 | ns |
| 122 | LABC-01-122 | 69.31 | 0.11 | ** | 2.82 | 0.56 | ns |
| 123 | LABC-01-123 | 88.85 | 0.14 | *** | 3.3 | 0.66 | ns |
| 124 | LABC-01-124 | 69.56 | 0.11 | ** | 3.23 | 0.65 | ns |
| 125 | LABC-01-125 | 75.89 | 0.12 | ** | 2.38 | 0.48 | ns |
| 126 | LABC-01-126 | 70.01 | 0.11 | ** | 3.83 | 0.77 | * |
| 127 | LABC-01-127 | 65.81 | 0.11 | ** | 3.62 | 0.72 | ns |
| 128 | LABC-01-128 | 49.83 | 0.08 | ns | 3.98 | 0.8 | * |
| 129 | LABC-01-129 | 80.18 | 0.13 | ** | 3.81 | 0.76 | * |
| 130 | LABC-01-130 | 75.02 | 0.12 | ** | 4.04 | 0.81 | * |
| 131 | LABC-01-131 | 69.32 | 0.11 | ** | 4.31 | 0.86 | * |
| 132 | LABC-01-132 | 77.23 | 0.12 | ** | 1.65 | 0.33 | ns |
| 133 | LABC-01-133 | 64.97 | 0.1 | ** | 2.15 | 0.43 | ns |
| 134 | LABC-01-134 | 82.68 | 0.13 | ** | 2.76 | 0.55 | ns |
| 135 | **LABC-01-135** | **31.8** | **0.05** | **ns** | **2.82** | **0.56** | **ns** |
| 136 | LABC-01-136 | 52.58 | 0.08 | ns | 4.85 | 0.97 | ** |
| 137 | LABC-01-137 | 65.52 | 0.11 | ** | 3.13 | 0.63 | ns |
| 138 | LABC-01-138 | 88.85 | 0.14 | *** | 3.13 | 0.63 | ns |
| 139 | LABC-01-139 | 83.46 | 0.13 | ** | 3.85 | 0.77 | * |
| 140 | LABC-01-140 | 76.34 | 0.12 | ** | 4.85 | 0.97 | ** |
| 141 | LABC-01-141 | 73.49 | 0.12 | ** | 2.08 | 0.42 | ns |
| 142 | LABC-01-142 | 82.82 | 0.13 | ** | 4.48 | 0.9 | ** |
| 143 | LABC-01-143 | 66.35 | 0.11 | ** | 2.82 | 0.56 | ns |
| 144 | LABC-01-144 | 60.53 | 0.1 | * | 2.9 | 0.58 | ns |
| 145 | LABC-01-145 | 95 | 0.15 | *** | 3.48 | 0.7 | ns |

| Entry | Genotype | A0 | SEM | Sig. | SB | SEM | Sig. |
|---|---|---|---|---|---|---|---|
| 146 | LABC-01-146 | 61.64 | 0.1 | * | 3.53 | 0.71 | ns |
| 147 | LABC-01-147 | 95 | 0.15 | *** | 4.07 | 0.81 | * |
| 148 | LABC-01-148 | 50.41 | 0.08 | ns | 2.08 | 0.42 | ns |
| 149 | LABC-01-149 | 51.53 | 0.08 | ns | 2.29 | 0.46 | ns |
| 150 | LABC-01-150 | 57.17 | 0.09 | * | 3.11 | 0.62 | ns |
| 151 | LABC-01-151 | 44.84 | 0.07 | ns | 3.11 | 0.62 | ns |
| 152 | LABC-01-152 | 66.42 | 0.11 | ** | 4.22 | 0.84 | * |
| 153 | LABC-01-153 | 77 | 0.12 | ** | 2.92 | 0.59 | ns |
| 154 | LABC-01-154 | 61.25 | 0.1 | * | 3.78 | 0.76 | * |
| 155 | **LABC-01-155** | **22.15** | **0.04** | ***** | **2.6** | **0.52** | **ns** |
| 156 | LABC-01-156 | 82.82 | 0.13 | ** | 2.24 | 0.45 | ns |
| 157 | LABC-01-157 | 63.07 | 0.1 | * | 4.58 | 0.92 | ** |
| 158 | LABC-01-158 | 57.85 | 0.09 | * | 3.17 | 0.64 | ns |
| 159 | LABC-01-159 | 91.54 | 0.15 | *** | 2.76 | 0.55 | ns |
| 160 | LABC-01-160 | 83.2 | 0.13 | ** | 4.31 | 0.86 | * |
| 161 | LABC-01-161 | 62.58 | 0.1 | * | 3.66 | 0.73 | ns |
| 162 | LABC-01-162 | 74.83 | 0.12 | ** | 2.88 | 0.58 | ns |
| 163 | LABC-01-163 | 51.22 | 0.08 | ns | 4.16 | 0.83 | * |
| 164 | LABC-01-164 | 43 | 0.07 | ns | 3.67 | 0.73 | ns |
| 165 | LABC-01-165 | 84.16 | 0.14 | *** | 4.79 | 0.96 | ** |
| 166 | LABC-01-166 | 63.19 | 0.1 | ** | 5.3 | 1.06 | ** |
| 167 | LABC-01-167 | 83 | 0.13 | ** | 2.82 | 0.56 | ns |
| 168 | LABC-01-168 | 88.01 | 0.14 | *** | 3.11 | 0.62 | ns |
| 169 | LABC-01-169 | 74.72 | 0.12 | ** | 3.67 | 0.73 | ns |
| 170 | LABC-01-170 | 95 | 0.15 | *** | 3.62 | 0.72 | ns |
| 171 | LABC-01-171 | 57.55 | 0.09 | * | 2.82 | 0.56 | ns |
| 172 | **LABC-01-172** | **31.69** | **0.05** | **ns** | **2.88** | **0.58** | **ns** |
| 173 | LABC-01-173 | 76.41 | 0.12 | ** | 2.76 | 0.55 | ns |
| 174 | LABC-01-174 | 79.98 | 0.13 | ** | 4.87 | 0.97 | ** |
| 175 | LABC-01-175 | 64.72 | 0.1 | ** | 3.67 | 0.73 | ns |
| 176 | LABC-01-176 | 90.62 | 0.15 | *** | 4.79 | 0.96 | ** |
| 177 | LABC-01-177 | 69.65 | 0.11 | ** | 3.74 | 0.75 | ns |
| 178 | LABC-01-178 | 67.76 | 0.11 | ** | 4.35 | 0.87 | * |
| 179 | LABC-01-179 | 44.32 | 0.07 | ns | 3.3 | 0.66 | ns |
| 180 | LABC-01-180 | 71.75 | 0.12 | ** | 3.3 | 0.66 | ns |
| 181 | LABC-01-181 | 51.39 | 0.08 | ns | 2.56 | 0.51 | ns |
| 182 | LABC-01-182 | 86.66 | 0.14 | *** | 4.64 | 0.93 | ** |
| 183 | LABC-01-183 | 83.2 | 0.13 | ** | 4.64 | 0.93 | ** |

| Entry | Genotype | A0 | SEM | Sig. | SB | SEM | Sig. |
|---|---|---|---|---|---|---|---|
| 184 | LABC-01-184 | 67.56 | 0.11 | ** | 3.42 | 0.68 | ns |
| 185 | LABC-01-185 | 51.98 | 0.08 | ns | 2.62 | 0.52 | ns |
| 186 | LABC-01-186 | 95 | 0.15 | *** | 4.76 | 0.95 | ** |
| 187 | LABC-01-187 | 83.31 | 0.13 | ** | 3.32 | 0.67 | ns |
| 188 | LABC-01-188 | 56.97 | 0.09 | * | 2.88 | 0.58 | ns |
| 189 | LABC-01-189 | 33.36 | 0.05 | ns | 3.94 | 0.79 | * |
| 190 | LABC-01-190 | 74.6 | 0.12 | ** | 4.51 | 0.9 | ** |
| 191 | LABC-01-191 | 71.59 | 0.12 | ** | 3.74 | 0.75 | ns |
| 192 | LABC-01-192 | 65.57 | 0.11 | ** | 3 | 0.6 | ns |
| 193 | LABC-01-193 | 56.13 | 0.09 | * | 4.38 | 0.88 | * |
| 194 | LABC-01-194 | 65.26 | 0.1 | ** | 3.04 | 0.61 | ns |
| 195 | LABC-01-195 | 84.16 | 0.14 | *** | 4.12 | 0.82 | * |
| 196 | LABC-01-196 | 31.15 | 0.05 | ns | 3.8 | 0.76 | * |
| 197 | LABC-01-197 | 72.43 | 0.12 | ** | 3.3 | 0.66 | ns |
| 198 | LABC-01-198 | 61.8 | 0.1 | * | 3.48 | 0.7 | ns |
| 199 | LABC-01-199 | 87.54 | 0.14 | *** | 4 | 0.8 | * |
| 200 | LABC-01-200 | 46.23 | 0.07 | ns | 3.91 | 0.78 | * |
| 201 | LABC-01-201 | 79 | 0.13 | ** | 4.25 | 0.85 | * |
| 202 | LABC-01-202 | 73.32 | 0.12 | ** | 4.85 | 0.97 | ** |
| 203 | LABC-01-203 | 65.34 | 0.11 | ** | 3.04 | 0.61 | ns |
| 204 | LABC-01-204 | 51.48 | 0.08 | ns | 3.56 | 0.71 | ns |
| 205 | LABC-01-205 | 74.65 | 0.12 | ** | 4.48 | 0.9 | ** |
| 206 | LABC-01-206 | 56.84 | 0.09 | * | 3.16 | 0.63 | ns |
| 207 | LABC-01-207 | 73.73 | 0.12 | ** | 4.31 | 0.86 | * |
| 208 | LABC-01-208 | 75.82 | 0.12 | ** | 4.16 | 0.83 | * |
| 209 | LABC-01-209 | 58.69 | 0.09 | * | 3.23 | 0.65 | ns |
| 210 | LABC-01-210 | 83.38 | 0.13 | ** | 3.04 | 0.61 | ns |
| 211 | LABC-01-211 | 71.94 | 0.12 | ** | 2.66 | 0.53 | ns |
| 212 | LABC-01-212 | 70.56 | 0.11 | ** | 2.56 | 0.51 | ns |
| 213 | LABC-01-213 | 86.8 | 0.14 | *** | 3.3 | 0.66 | ns |
| 214 | LABC-01-214 | 65.41 | 0.11 | ** | 3.11 | 0.62 | ns |
| 215 | LABC-01-215 | 62.27 | 0.1 | * | 3.56 | 0.71 | ns |
| 216 | LABC-01-216 | 68.79 | 0.11 | ** | 2.9 | 0.58 | ns |
| 217 | LABC-01-217 | 82.38 | 0.13 | ** | 3.27 | 0.65 | ns |

SEM, Standard error of means; Sig., Significance level in relation to LR-59-81; *, ** and *** indicate significance difference at $P \leq 0.05$, 0.01, 0.001, respectively; ns, not-significant. A0 Levene's test (F-value=2.12, $P<0.0001$), SB Levene's test (F-value =1.92, $P<0.0001$).
