## Supplementary Figure S1 for "An advanced lentil backcross population developed from a cross between *Lens culinaris* × *L. ervoides* for future disease resistance and genomic studies"

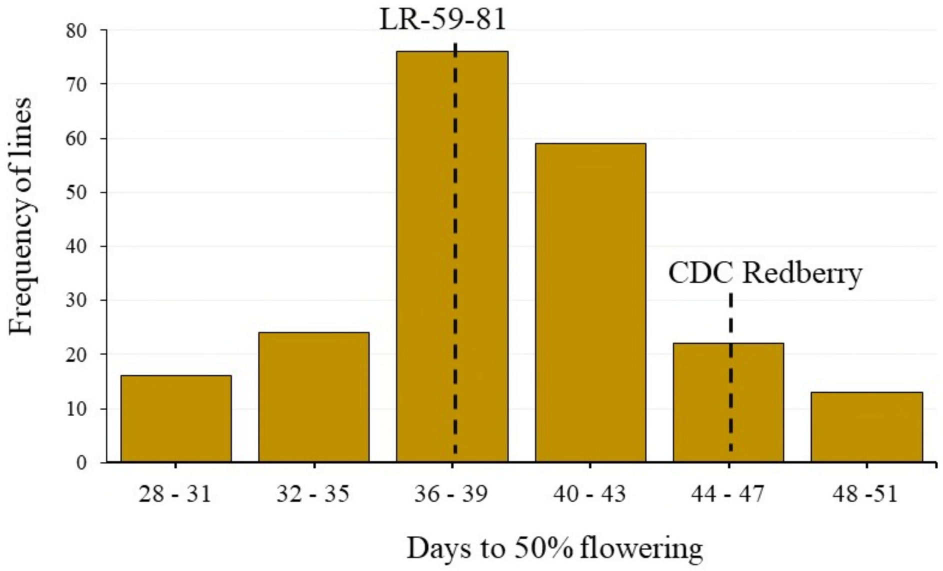


**Supplemental Figure S1.** Frequency distribution of days to 50% flowering in the LABC-01 population at $BC_2F_{3:4}$ generation. The data was observed under growth chamber conditions.
